# Cross-Site Comparison of Ribosomal Depletion Kits for Illumina RNAseq Library Construction

**DOI:** 10.1101/212639

**Authors:** Zachary T. Herbert, Jamie P. Kershner, Vincent L. Butty, Jyothi Thimmapuram, Sulbha Choudhari, Yuriy O. Alekseyev, Jun Fan, Jessica W. Podnar, Edward Wilcox, Jenny Gipson, Allison Gillaspy, Kristen Jepsen, Sandra Splinter BonDurant, Krystalynne Morris, Maura Berkeley, Ashley LeClerc, Stephen D. Simpson, Gary Sommerville, Leslie Grimmett, Marie Adams, Stuart S. Levine

## Abstract

Ribosomal RNA (rRNA) comprises at least 90% of total RNA extracted from mammalian tissue or cell line samples. Informative transcriptional profiling using massively parallel sequencing technologies requires either enrichment of mature poly-adenylated transcripts or targeted depletion of the rRNA fraction. The latter method is of particular interest because it is compatible with degraded samples such as those extracted from FFPE and also captures transcripts that are not poly-adenylated such as some non-coding RNAs. Here we provide a cross-site study that evaluates the performance of ribosomal RNA removal kits from Illumina, Takara/Clontech, Kapa Biosystems, Lexogen, New England Biolabs and Qiagen on intact and degraded RNA samples. We find that all of the kits are capable of performing significant ribosomal depletion, though there are differences in their ease of use. All kits were able to remove ribosomal RNA to below 20% with intact RNA and identify ∼14,000 protein coding genes from the Universal Human Reference RNA sample at >1FPKM. Analysis of differentially detected genes between kits suggests that transcript length may be a key factor in library production efficiency. These results provide a roadmap for labs on the strengths of each of these methods and how best to utilize them.

## INTRODUCTION

Ribosomal depletion is a critical method in transcriptomics that allows for efficient detection of functionally relevant coding as well as non-coding transcripts through removal of highly abundant rRNA species. Use of oligo dT primer to capture the polyadenylated 3’ end of the transcripts and isolate mRNA is routine in many RNA sequencing preparations; however this method lacks the ability to handle degraded samples where most of the RNA is separated from the 3’ tail, or to isolate non-polyadenylated transcripts such as lncRNAs. Ribosomal removal methods address these issues by directly depleting the rRNA while leaving other transcripts intact. This technique is widely utilized and is a basic component of many large datasets (Cui et al. 2010; Zhao et al. 2014; Guo et al. 2015).

The current generation of rRNA removal kits employs three distinct strategies to deplete these transcripts. In the first method, rRNA is captured by complimentary oligonucleotides that are coupled to paramagnetic beads, after which the bound rRNA is precipitated and removed from the reaction. Kits utilizing this method include Illumina’s RiboZero, Qiagen GeneRead rRNA depletion, and Lexogen RiboCop. The second method uses an alternative strategy, hybridizing the rRNA to DNA oligos and degrading the RNA:DNA hybrids using RNAseH. These kits include NEBNext rRNA depletion, Kapa RiboErase, and Takara/Clontech’s RiboGone. A third method that is specifically aimed at low-input samples using the Takara/Clontech SMARTer Pico kit, targets the ribosomal RNA sequences after conversion to cDNA and library prep using the ZapR enzyme (CLONTECH WEBSITE). Additional high abundance transcripts such as globin and mitochondrial RNA (mtRNA) can be targeted by each method. Despite the availability of many different kits utilizing these methods, the efficiency of rRNA removal and possible off-target effects of these different methodologies on the resulting RNAseq data remains unclear.

With an increasing number of ribosomal RNA depletion kits available, understanding the relative strengths of these methods is critical for improving experimental design. To address this challenge, we have conducted a cross-site study comparing seven rRNA depletion kits against a standard sample both as intact and degraded RNA. We find that about half of the kits are likely to require significant care in implementation and note that the Lexogen RiboCop and Qiagen GeneRead kits worked poorly with heavily degraded samples. The different kits also appear to be affected by relative lengths of the transcripts as well as the degradation of the input RNA. These results suggest that different methodologies may be appropriate depending on the experimental question and quality of input material.

## RESULTS

In order to better understand the strengths of the different ribosomal depletion methodologies available, we utilized a set of controlled samples that could be broadly distributed in order to provide a consistent biological background for each kit and site (Figure 1). All experiments utilized the well characterized Universal Human Reference RNA (UHR) from Agilent, either in its intact state, or following heat degradation (Figure S1). Two control spike-ins were added to this sample, the Lexogen Spike-In RNA Variant Controls (SIRVs) which were added before degradation and co-degraded with the sample, and the External RNA Controls Consortium (ERCC) from Ambion, which was added after degradation and thus remained intact and served as an additional control. For each experiment, 100ng of RNA input was used with the exception of the Takara/Clontech SMARTer pico kit which used 1ng input as recommended by the manufacturer.

**Figure 1:**
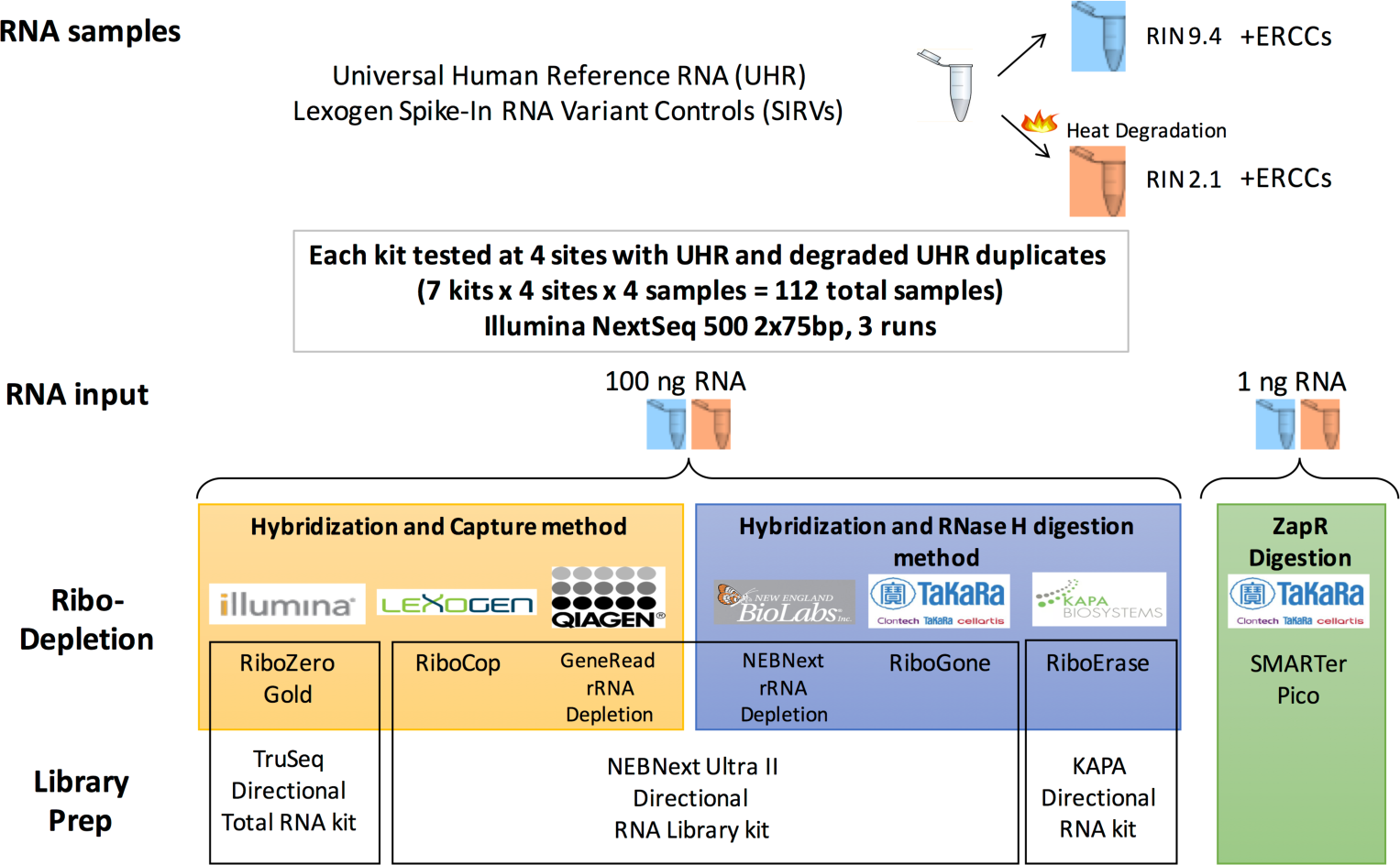
Design of the ribosomal depletion study. Schematic of the sample processing is shown. A single sample of UHR RNA with SIRV spike-ins was kept intact or heat degraded followed by addition of the ERCC spike-in. The two samples were then distributed to the participating sites where they were run as technical duplicates for each kit.

The study tested seven rRNA depletion kits (Figure 1), each tested at four sites. The kits tested include Illumina RiboZero Gold (RZ), Lexogen RiboCop (LX), Qiagen GeneRead rRNA Depletion (Q), all of which use bead capture for ribosomal depletion, the New England Biolabs NEBNext rRNA Depletion (NE), Kapa RiboErase (K), and Takara/Clontech Ribogone (CR) kits that are based on RNAseH degradation of the rRNA, and SMARTer Pico (CZ) which uses the ZapR enzyme to remove rRNA after library prep. The Lexogen, Qiagen, Ribogone, and NEBNext kits all utilized the NEB Next Ultra II directional library generation kit to convert the RNA to Illumina libraries while the other kits used RNA library generation kits from the manufacturer of the depletion chemistry. A total of 11 sites participated in the study with each site handling no more than four kits. Sites were selected from genomic core facilities that are members of the Association of Biomedical Research Facilities (ABRF) who routinely preform library preparation for academic labs. For each vendor, a consultation conference call was held between the vendor and the participating sites to review the protocol in detail before the experiment was performed with the goal of standardizing and clarifying the protocol, thereby minimizing the chance for confusion about the methodology. Technical duplicates of both the degraded and intact RNA were run at each site for each kit. The total 106 samples after dropouts (see supplemental text) were pooled and sequenced on three NextSeq500 runs at a single site to eliminate bias due to sequencing.

The different methodologies were first evaluated for their ability to perform their primary objective, the removal of rRNA from the samples before sequencing. rRNA reads in each sample were identified by aligning to known rRNA sequences using BWA (Li and Durbin 2009). A cutoff of 50% nuclear rRNA was chosen to indicate ribosomal depletion failure. Illumina’s RiboZero Gold kit showed ∼5% rRNA with the intact sample at all sites (excluding a single point failure) but slightly higher rRNA fractions for the degraded sample. This kit was used as a baseline for the other kits due to its long-standing reputation in the sequencing community (Figure 2a). The other kits that used the capture method for depletion were less consistent than RiboZero Gold with one failed site for Lexogen RiboCop and three failed sites for the Qiagen GeneRead rRNA depletion kit. For both the Lexogen and Qiagen kits, the intact samples performed significantly better than the degraded sample and caution should be used when using these kits on highly degraded RNA.

**Figure 2:**
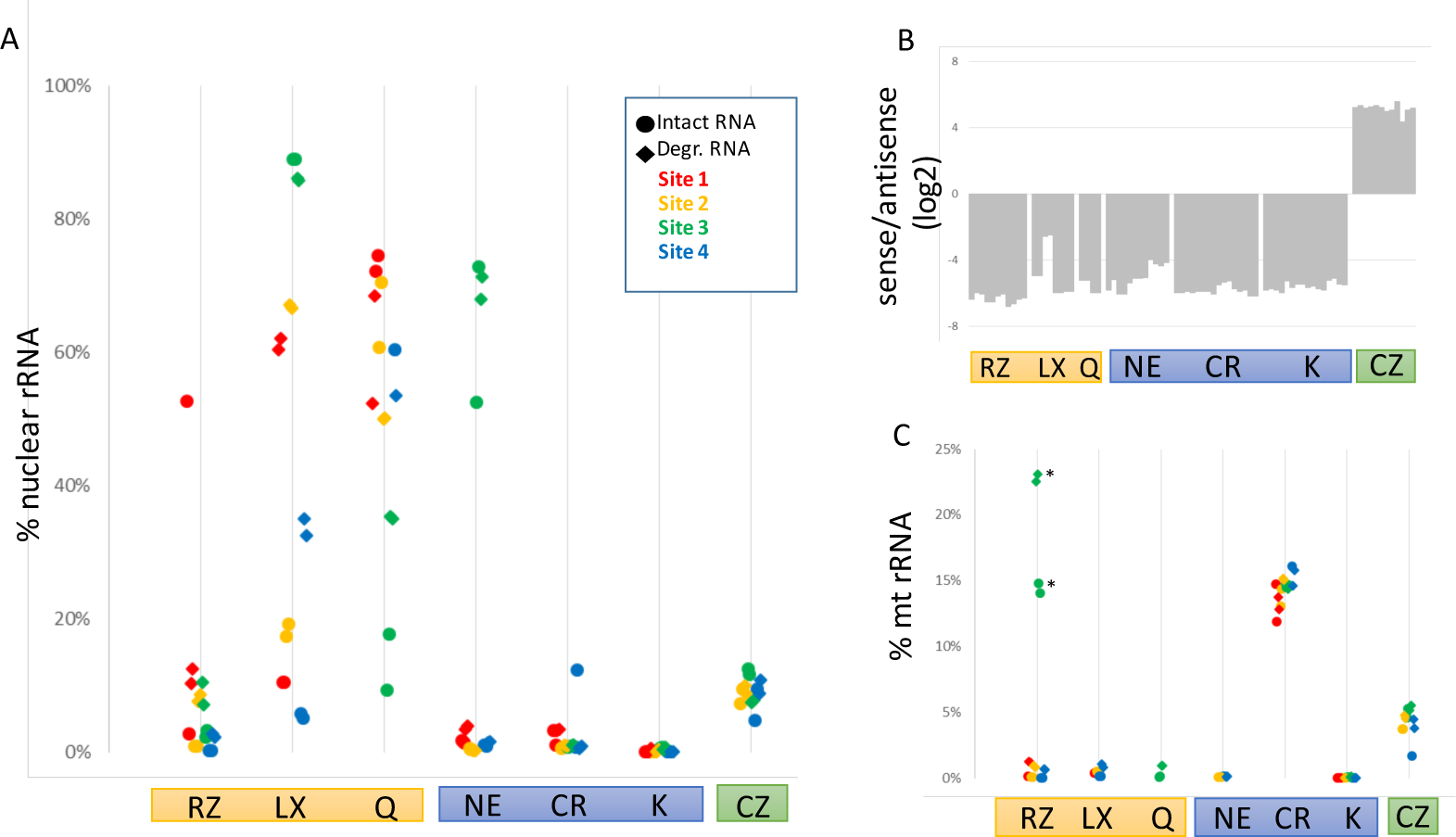
Properties of the rRNA depleted libraries. A) Fraction of reads mapping to nuclear rRNA shown. Site number indicated by color. Intact samples are shown as circles, degraded samples are shown as diamonds. Kit abbreviations: RZ=RiboZero Gold, LX=Lexogen RiboCop, Q=Qiagen GeneRead rRNA Depletion, NE=NEBNext rRNA Depletion, K=Kapa RiboErase, CR=Clontech Ribogone, CZ=SMARTer Pico total RNA. B) Reads were mapped to exons in UCSC known gene and scored based on strand of alignment. C) Fraction of reads mapping to mt rRNA shown as in A. *-RiboZero site 3 used standard RiboZero instead of RiboZero Gold.

By comparison, the kits that degraded the rRNAs by either RNase H treatment or using ZapR showed more consistent results. Excluding single sites that failed with the NEBNext Ultra rRNA and SMARTer Pico kits, those two kits as well as the Takara/Clontech RiboGone and Kapa RiboErase kits performed very well with no differences observed between intact and degraded RNA. The RNaseH methods all showed very low rRNA fractions overall with the noted exception to the NEB kit. The SMARTer Pico kit had a slightly higher rRNA level, similar to that observed with Illumina Ribozero Gold degraded samples.

For those samples with successful rRNA depletion, we next ascertained the quality of the RNA sequencing data. Samples with greater than 50% rRNA were excluded from further analysis to eliminate artifacts that may be caused by improper implementation of the protocol. All of the kits showed strong strand bias as expected by the protocols (Figure 2b). Notably, SMARTer Pico reads mapped to the sense strand, which is the opposite strand from the other methods. While this is expected, care should be taken in adapting existing informatics pipelines to this kit. Differences were observed among the kits in how they handled mtRNAs. These were a major contaminant in the Clontech kits, particularly the RiboGone method that only targets the 12s mtRNA and not the 16s mtRNA. The other methods that addressed all mtRNA reads, such as Illumina Ribozero Gold, significantly reduced the fraction of reads from mtRNAs (figure 2c). This is especially noticeable in the RiboZero samples where site3 utilized a standard Human/Mouse/Rat kit instead of the RiboZero Gold.

Looking at the non-rRNA reads in each sample, the vast majority align to protein coding genes based on the ENSEMBL annotation. All samples show >60% protein coding with most over 80% and the Clontech RiboGone kit having the largest fraction mapping (Figure 3a). Most of the samples identified ∼14,000 genes expressed at greater than one RPKM and ∼16,000 at over 0.1 RPKM (Figure 3b). A single site using the Takara/Clontech SMARTer Pico kit did show a somewhat reduced number of genes, which was associated with a lower library complexity observed from that site. Antisense mapped reads and reads mapping to the signal recognition particle RNAs (SRPs) were the most variable aspect of each sample though the source of these differences were unclear as they varied widely from site to site.

**Figure 3:**
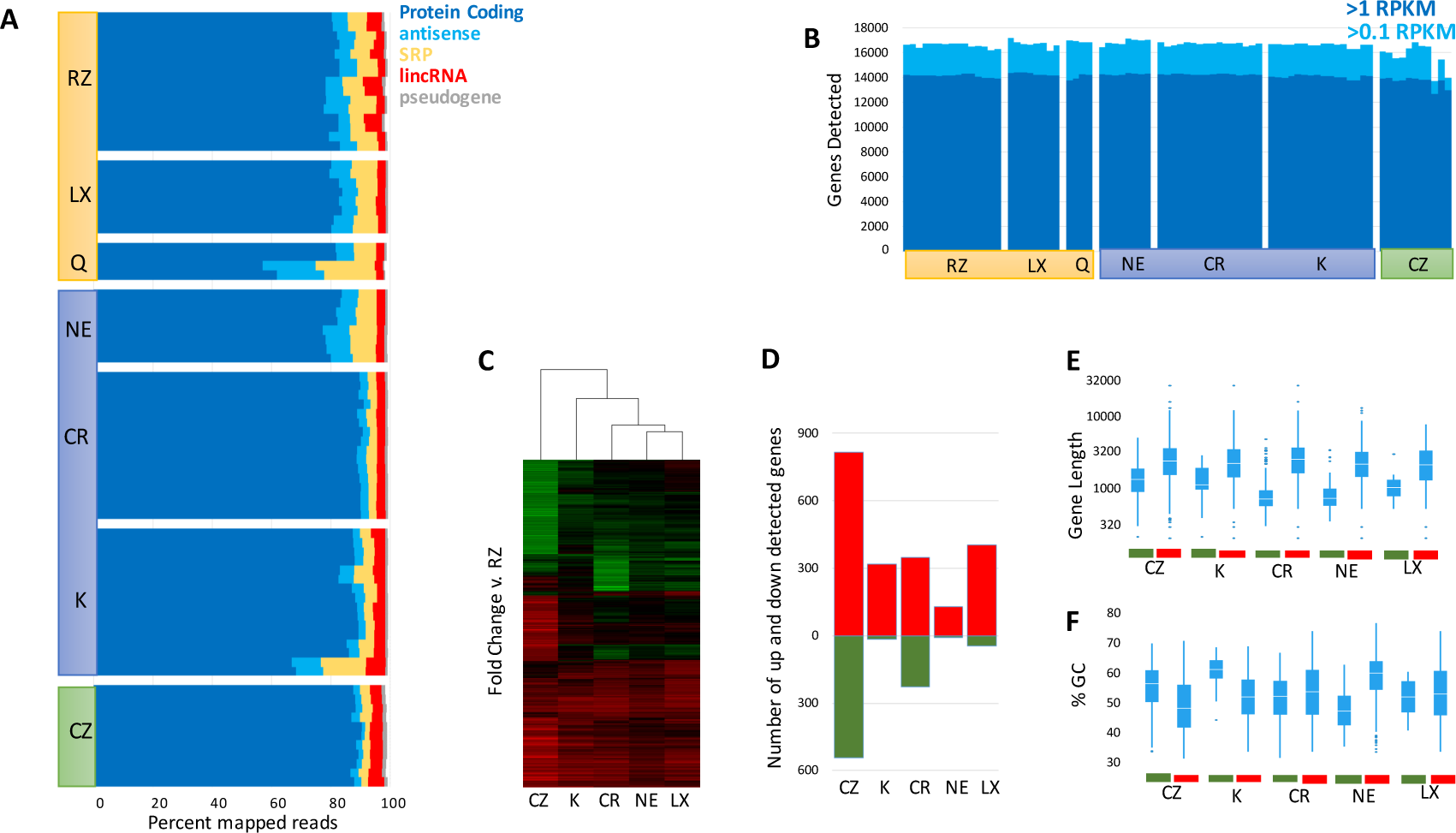
Protein Coding Gene Detection in rRNA Depleted Libraries. A) Non-mtRNA reads were mapped to the ENSENBL annotation and grouped by transcript type. Fraction of reads associated with transcript types >1% shown. Data sets ordered by site then intact/degraded status within each kit top to bottom. B) Number of genes detected at >1RPKM (dark blue) and >0.1RPKM (light blue) shown for each replicate. Genes ordered the same as in A but left to right. C) Changes in RNA detection compared to Illumina RiboZero. Hierarchical clustering [X] genes with fold changes >2 and Benjamini corrected p-values < 0.001 are shown (union of all comparisons). D) Count of genes with fold changes >2 and Benjamini corrected p-values < 0.001 are shown for each kit as compared to Illumina RiboZero. Increased detection shown in red, decreased detection shown in green. E) Distribution of read lengths for transcripts detected at higher (red) or lower (green) rate relative to RiboZero. F) Distribution of GC% for transcripts detected at higher (red) or lower (green) rate relative to RiboZero.

While the total number of protein coding genes detected was quite similar, many genes appear to be detected at significantly different rates. To better understand this observation, we performed differential gene expression analysis on the intact RNA samples that passed our QC metrics, comparing each preparation back to Illumina RiboZero Gold (Figure 3c). The Qiagen rRNA depletion kit was excluded as only two replicates passed these criteria. Hierarchical clustering of the differentially detected genes, clusters the kits first by their RNAseq library prep methodology, with all three kits prepared using the NEB Ultra II Directional RNA Library kit clustering together, followed by the Kapa RiboErase kit and finally the low input Takara/Clontech SMARTer pico. Generally, several hundred genes could be easily observed as differentially detected between each of the kits and Illumina’s RiboZero (fold changes >2, Benjamini corrected p-values < 0.001, Figure 3d). Testing the physical properties of these differentially detected genes found that gene length appears to be a large contributor to the direction of the bias with shorter transcripts better detected by RiboZero and longer transcripts better detected by the other kits (Figure 3e). Many of the most variably detected genes across the data set are quite small, such as mitochondrial proteins and ribosomal proteins (Figure S2). The libraries themselves did not display any particular size bias with the RiboZero samples having an average length distribution similar to the other kits (Figure S3). Bias in GC percentage was also observed in a few samples, with the Kapa RiboErase kit having the strongest bias against high GC transcripts (though the underlying gene list is quite small, Figure 3f).

Ribosomal depletion is a key methodology used in studying noncoding RNAs as many of these transcripts lack polyadenylation sites (Ulitsky and Bartel 2013). We focused on lincRNAs (long intervening noncoding RNAs), one type of noncoding RNAs, since they do not overlap with any protein coding or other long non-coding RNA genes. Using the ENSEMBL annotation, approximately 4% of the non rRNA reads map to lincRNAs using RiboZero (Figure 4a). Similar numbers are observed with the SMARTer Pico kit and Kapa RiboErase. The other kits, all of which were prepared with NEB Ultra II directional RNA library kit, show less than 3% of reads mapping to lincRNAs. This global decrease in number of lincRNA reads appears to reflect a general decrease in the number of mapping reads rather than a specific bias against a subset of lincRNAs. Two lines of evidence support this conclusion. First, the number of lincRNAs detected at >=0.01 RPM remains ∼3500 for all of the different kits tested and no bias is seen against the NEB prepped kits (Figure 4b). Second, while the majority of lincRNAs detected can be assigned to 4 specific lincRNAs (MALAT1, SNORD3A, RNRP and NEAT1), the remaining fraction remains constant among the different kits, suggesting a global decrease in mapping (Figure 4c). The precise set of lincRNAs detected varies somewhat but the core of 3200 lincRNAs are detected by all of the kits (Figure 4d).

**Figure 4:**
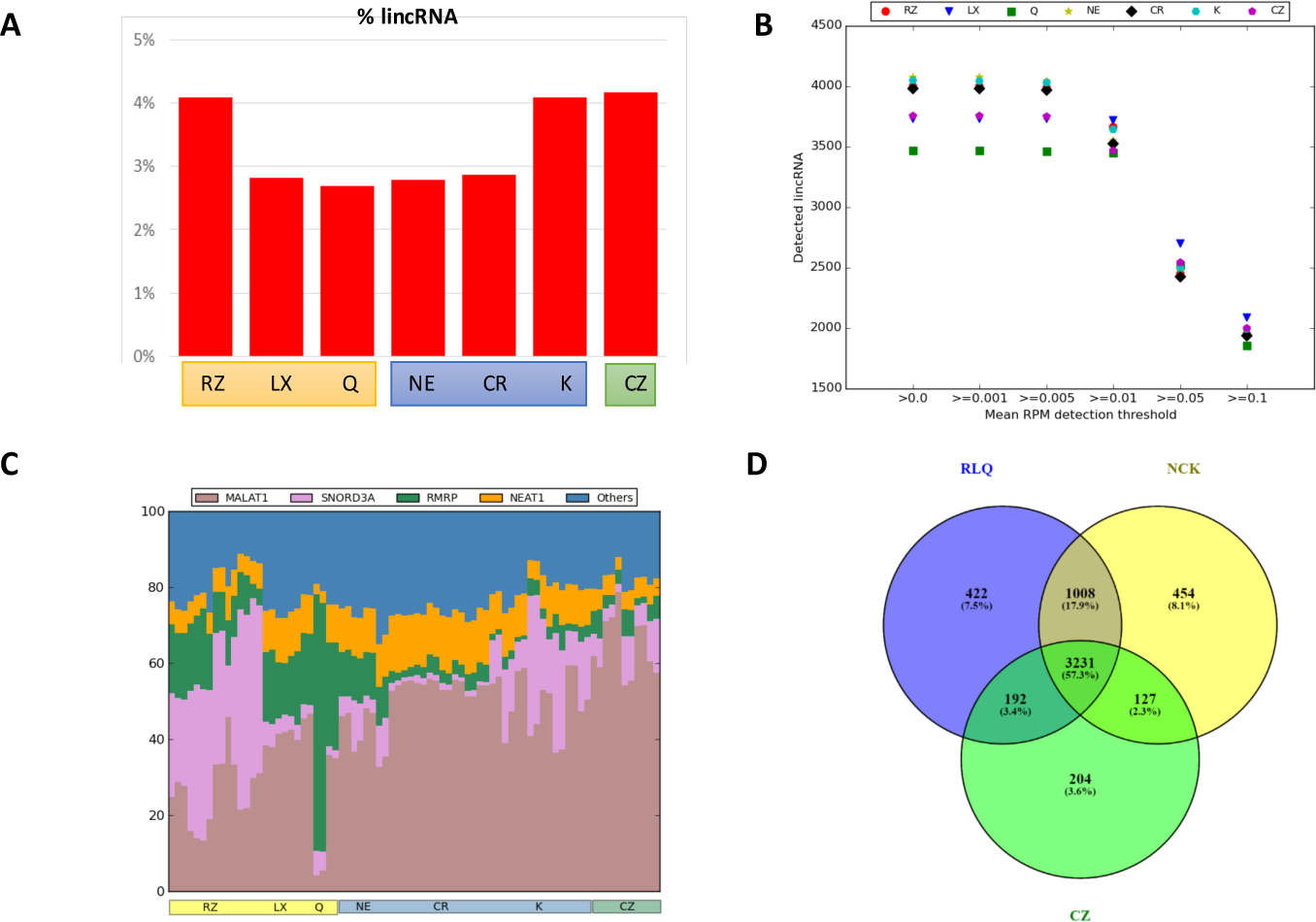
LincRNA Detection in rRNA Depleted Libraries. A) Mean fraction of non-rRNA reads assigned to lincRNAs based on the ENSEMBL annotation. B) Number of lincRNAs detected over specified RPM levels. C) Fraction of lincRNA mapped reads assigned to the top 4 lincRNAs detected for each sample. Data sets ordered by site then intact/degraded status within each kit left to right. Average RPM counts for each lincRNA was calculated and the top four lincRNAs were shown keeping remaining lincRNAs in ‘Others’ category. D) Overlap of lincRNAs detected by the three core library prep methodologies: ribosomal pulldown (RLQ), RNAse H (NCK), and ZapR (CZ). Average RPM counts for each lincRNA for all samples in each of the three core library methods (RLQ = RZ, LX, Q; NCK = NE, CR, K; CZ = CZ) was calculated and lincRNAs with average RPM > 0 were compared among the methods.

While comparison of different detection rates of mRNAs in UHR can point to possible differences between the kits’ chemistries, the spike-in controls provide an absolute metric to evaluate their effectiveness. Both SIRV (co-degraded with the RNA) and ERCC (not degraded) control spike-ins were added to the sample before library preparation and should give an unbiased look at the behavior of the different kits. As an initial test, we examined the ratio between the two spike-in types to model the impact of degradation on efficiency of library formation. Importantly, the kits do treat degraded RNA differently in their protocols and the not degraded ERCCs in the degraded samples were processed along the path proscribed for the bulk RNA, suggesting they are under-fragmented relative to the bulk population. Examining the ratios between the ERCC and SIRVs, we find that all the intact samples show ∼60% SIRV reads (Figure 5a). By comparison, the degraded samples show significant bias between the ERCCs and SIRV, generally favoring the intact ERCC. The RiboZero Gold and Kapa RiboErase kits show the least bias based on degradation, while the kits using the NEB stranded RNAseq kits showed a bias against shorter RNA fragments, which is similar to what was observed for protein coding genes (Figure 3e). The SMARTer Pico kit is biased against the intact ERCC spike-ins in the degraded sample, which is likely due to not pre-degrading the ERCCs in the context of the degraded total RNA, emphasizing the importance of this step. Inserting not degraded spike-in controls into variably degraded experimental samples may confound the ability of these spike-ins to serve as a normalization tool, as previously observed (Jiang et al. 2011; Risso et al. 2014).

**Figure 5:**
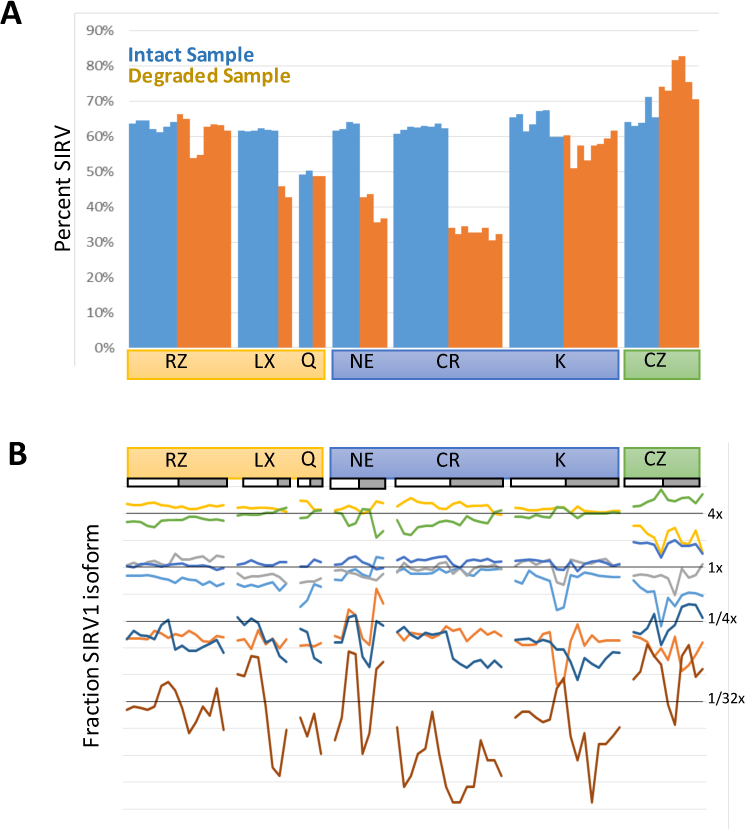
Effect of rRNA Depletion Chemistry on Spike-In Controls. Effect of degradation of SIRVs on the ratio of SIRV reads to ERCC reads in each replicate. Percent of reads mapping to SIRVs out of total reads mapping to Spike-ins in shown. Data sets ordered by intact/degraded status followed by site within each kit left to right. B) Relative ratio of reads mapping to SIRV1. Fraction of reads mapping to each isoform of SIRV1 are shown for each replicate. Expected fractions shown as dark horizontal lines. Light horizontal lines show 2-fold changes in fraction observed (log scale). Each SIRV1 isoform is shown in a different color. Replicates ordered as in A.

Distinct transcripts are present at defined ratios in the spike-in control allowing direct visualization of over and under-representation of transcripts. Within the SIRV spike-ins, the distinct transcripts were largely at equal ratios between kits and sites, though deviation from the expected values is observed (Figure 5b). The SMARTer Pico kit was particularly susceptible to variation, possibly due to the low total input (1:100^th^ of the other kits), and some transcripts (e.g. purple at 1/4x) show loss of signal in the degraded sample. By comparison, the ERCC spike-ins showed significantly more variability across sites, even within the intact RNA samples (Figure S3). Overall, the Lexogen and Takara/Clontech RiboGone kits generally had the most consistent and even performance on both the ERCC and SIRV spike-in controls across their test sites for intact samples.

## DISCUSSION

rRNA depletion methodologies have expanded significantly in the last year. While differences exist between the rRNA depletion chemistries tested, all of the kits tested were able to successfully remove a significant amount of the rRNA in library preparations. The bead depletion chemistries were the most challenging to consistently implement successfully and struggled to remove rRNAs in the degraded RNA samples. All of the kits, including the low-input ZapR based kit from Takara/Clontech, detected ∼14,000 transcripts at >1 RPKM.

With the broad success of the different chemistries, other aspects of library preparation, such as cost and ease of use, can be considered. Participating sites were surveyed to collect feedback regarding ease of use and previous experience with each kit that was tested (Table 1). While all sites were familiar with the RiboZero depletion kit, the majority of sites had no prior experience with the other kits. The participating sites generally reported high comfort levels with all of the kits. Interestingly, the comfort level with the different kits did not correlate with success as the sites with the lowest comfort levels for the NEB kit and ZapR based kit from Takara/Clontech both showed very good performance. This may be due to the robustness of the specific methods or the quality of the written protocols.

While we did observe differences in the efficiency of library preparation, analytics remains a key caveat. For this study we used STAR/RSEM/DESEQ (Dobin et al. 2013; Love et al. 2014; Li and Dewey 2011) for the analysis of the transcript levels, but different informatics tools may have more or less ability to handle the variations between the different chemistries and to model the spike-in controls. The combination of defined control samples with single transcript and spliced spike-ins provides an opportunity to use this data in the evaluation of different algorithmic approaches without overfitting to a single site or single type of chemistry. The number of RNAseq algorithms continues to multiply and finding the most appropriate methodology remains challenging. We believe this data set will provide a unique opportunity to better characterize the strengths and challenges of not only the depletion chemistries, but the RNAseq analysis algorithms as well.

## METHODS

### Input RNA preparation

Intact and degraded input RNA was prepared and aliquoted at a single site. The Universal Human Reference RNA (Agilent) was diluted to 500 ng/ul in 200ul of RNase-free water and 3.94ul of the Spike-in RNA Variant Control E2 Mix (Lexogen) were added. The sample was split into two aliquots, one of which was then heated at 94° C on an Eppendorf™Thermomixer for 1 hour and 27 minutes. 1ul of ERCC RNA Spike-In Mix 1 (ThermoFisher Scientific) was added to both the intact and degraded samples before running on a Bioanalyzer 2100 RNA Nano chip (Agilent) (Figure S1). The final intact and degraded RNA samples were then diluted to 25 ng/uL and were distributed to each site on dry ice for rRNA-depletion and library preparation.

### Site Selection and Index Allocation

Eleven genomics core facilities were selected from among the members of the Association of Biomedical Research Facilities membership. All of these cores routinely perform Illumina library preparation for laboratories at their institutes and each site prepared between one and four library types. Kits were assigned to each site to minimize overlapping sets with the exception of the Takara/Clontech and Illumina kits. All four Takara/Clontech sites preformed both the Ribogone (CR) and SMARTer Pico (CZ) kit to minimize shipping costs. Similarly, Illumina RiboZero (RZ) sites were selected to minimize shipping of the donated reagents. Indices were assigned by the group to prevent overlapping among libraries.

### rRNA Depletion and Library Construction

Each site performed rRNA depletion and subsequent library prep following the vendor protocols (Table S1). Conference calls were held with the vendors and recommended deviations from the protocols resulting from those meetings are outlined in the Supplementary methods. Input RNA concentrations, fragmentation conditions and PCR cycles for intact and degraded RNA samples for each kit were discussed with the vendors and can be found in Table S2 for each kit. Completed libraries were quantified by Qubit or equivalent and run on a Bioanalyzer or equivalent for size determination. Libraries were pooled and sent to a single site for final quantification by Qubit fluorometer (ThermoFisher Scientific), TapeStation 2200 (Agilent), and RT-qPCR using the Kapa Biosystems Illumina library quantification kit. Libraries were pooled for each run on a NextSeq 500.

### Sequencing

Sites were instructed to make an equimolar pool of libraries from each kit using site-specific quantification and pooling SOPs and return each pool along with individual un-pooled libraries to the designated sequencing site. The sequencing site quantified each pool by Qubit fluorometer, Agilent TapeStation, and qPCR using the Kapa Illumina quant assay. Library pools were multiplexed and sequenced over three high output paired-end 75bp runs on the Illumina NextSeq 500 to achieve sufficient read depth for analysis (See Supplemental Methods).

### Alignment and Quality Control

Reads were aligned against hg19 using bwa-mem v. 0.7.12-r1039 with flags −t 16 −f (Li and Durbin 2009). Mapping rates, fraction of multiply-mapping reads, strandedness (Figure 2B), number of unique 20-mers at the 5’ end of the reads, insert size distributions (Figure S3) and fraction of nuclear ribosomal RNAs (Figure 2A) were calculated using dedicated perl scripts and bedtools v. 2.25.0 (Quinlan and Hall 2010). In addition, each resulting bam file was randomly down-sampled to one million aligned reads and read density across genomic features were estimated for RNA-Seq-specific quality control metrics. Samples with <10 reads or >50% rRNA were eliminated from additional analysis and were not included in later sequencing pools. Sample reads from all runs were concatenated before final RNA analysis.

### RNA-Seq mapping and quantitation

Reads were aligned against hg19 / ENSEMBL 75 annotation using STAR v. 2.5.1b (Dobin et al. 2013) with the following flags -runThreadN 8 --runMode alignReads --outFilterType BySJout --outFilterMultimapNmax 20 --alignSJoverhangMin 8 --alignSJDBoverhangMin 1 --outFilterMismatchNmax 999 --alignIntronMin 10 --alignIntronMax 1000000 --alignMatesGapMax 1000000 --outSAMtype BAM SortedByCoordinate-quantMode TranscriptomeSAM with --genomeDir pointing to a low-memory footprint, 75nt-junction hg19 STAR suffix array. Gene expression was quantitated using RSEM v. 1.2.30 [9] with the following flags for all libraries: rsem-calculate --expression-paired-end --calc-pme --alignments -p 8 --forward-prob 0 against an annotation matching the STAR SA reference, with the exception of the positively-stranded CZ libraries, for which --forward-prob was set to 1. Posterior mean estimates (pme) of counts were retrieved, ribosomal RNA counts removed and expression in reads per kilobase of modeled exon per million mapped reads (RPKM, Figure 3B, S2) or transcripts per million (TPM, Figure 3A) were computed on the remaining count matrices. Similar scripts and pipelines were used for ERCC and SIRV mapping and quantitation. For both ERCCs and SIRVs, counts were retrieved from the RSEM gene output, as well as fractional isoform usage for SIRVs.

### Differential representation analysis

Libraries and kits were compared against RiboZero samples using a standard differential expression framework. Briefly, differential expression was performed in the R statistical environment (R v. 3.2.3) using Bioconductor’s DESeq2 package on the protein-coding genes only (Love et al. 2014). Dataset parameters were estimated using the estimateSizeFactors(), and estimateDispersions() functions, and differential expression based on a negative binomial distribution/Wald test was performed using nbinomWaldtest() (all packaged into the DESeq() function), using the kit type as a contrast. Fold-changes, p-values and Benjamin-Hochberg-adjusted p-values (BH) were reported for each gene. Genes with BH<0.001 and absolute fold-changes greater than 2 were considered for downstream analyses (Figure 3C-D).

### lincRNA analysis

After removing lincRNAs that were not assigned to chromosomes, 7251 lincRNAs were identified from the ENSEMBL annotation for further analysis. The lincRNAs with mean RPM >=0.01 were retained for each kit. To examine the distribution of different lincRNAs in each of the samples, the percentage RPM was calculated. Except the top 4 lincRNAs, rest of the lincRNAs were classified as ‘Others’ category. The figures were generated using a custom python script. Based on the <u>ribo</u>-depletion methods, the seven kits were grouped into three sets: RLQ (includes RZ, LX, and Q), NCK (includes NE, CR, K) and CZ. The lincRNAs detected in three sets were compared using an Euler diagram created using Venny (http://bioinfogp.cnb.csic.es/tools/venny/).

### Genomic features analysis and visualization

Gene lengths were retrieved from RSEM outputs. GC content was calculated on the longest annotated isoform of each protein-coding gene. Box-plots were generated using Spotfire (Tibco) and TreeView (Figure 3E-F).

## ACCESSION NUMBERS

The NCBI Gene Expression Omnibus accession number for the sequencing data reported is GSE100127.

**Table 1:** User Experience with Different rRNA Depletion Chemistries. Study participants were surveyed for their opinions on the ease of use of each kit they tested.

| Kit | Site | Used kit before | Comfort level* |
| --- | --- | --- | --- |
| RZ | 1 | Yes | 3-4 |
|  | 2 | Yes | 5 |
|  | 3 | Yes | 5 |
|  | 4 | Yes | 5 |
| LX | 1 | No | 5 |
|  | 2 | No | 5 |
|  | 3 | No | 3 |
|  | 4 | No | 3 |
| Q | 1 | No | 5 |
|  | 2 | No | 4 |
|  | 3 | No | 4 |
|  | 4 | No | 5 |
| NE | 1 | No | 2-3 |
|  | 2 | Yes | 4 |
|  | 3 | No | 4 |
|  | 4 | No | 5 |
| CR | 1 | No | 4 |
|  | 2 | No | 4 |
|  | 3 | No | 3-4 |
|  | 4 | No | 5 |
| K | 1 | No | 4 |
|  | 2 | Yes | 3-4 |
|  | 3 | No | 5 |
|  | 4 | No | 3-4 |
| CZ | 1 | No | 4 |
|  | 2 | No | 4 |
|  | 3 | Yes | 3-4 |
|  | 4 | No | 2 |

\*On a scale of 1-5 with 1 being not at all comfortable and 5 being very comfortable

## ACKNOWLEDGEMENTS

This study is a product of the DNA Sequencing Research Group (DSRG) of the Association of Biomedical Research Facilities (ABRF, www.abrf.org). The authors are grateful to the participating vendors: Illumina, New England Biolabs, Lexogen, Takara/Clontech, Qiagen and Kapa/Roche who donated library preparation reagents, spike-in controls and sequencing reagents as well as to the ABRF executive board for funding. We also thank the many member labs within the ABRF who volunteered to test kits. ZTH JPK JT YA, JF, JWP, MA, --This work is supported by NCI award P30-CA14051 (VB, SSL), NIEHS award P30-ES002109 (VB, SSL) and the Harvard University Center for Aids Research NIH award P30 AI060354 (ZTH,MB,GS,LG).

## REFERENCES

Cui, P., Q. Lin, F. Ding, C. Xin, W. Gong, L. Zhang, J. Geng, B. Zhang, X. Yu, J. Yang, S. Hu and J. Yu. 2010. A comparison between ribo-minus RNA-sequencing and polyA-selected RNA-sequencing. Genomics. 96(5): 259–265.

Dobin, A., C. A. Davis, F. Schlesinger, J. Drenkow, C. Zaleski, S. Jha, P. Batut, M. Chaisson and T. R. Gingeras. 2013. STAR: ultrafast universal RNA-seq aligner. Bioinformatics 29(1): 15–21.

Guo, Y., S. Zhao, Q. Sheng, M. Guo, B. Lehmann, J. Pietenpol, D. C. Samuels and Y. Shyr. 2015. RNAseq by Total RNA Library Identifies Additional RNAs Compared to Poly(A) RNA Library. Biomed Res Int. 2015: 862130.

Jiang, L., F. Schlesinger, C. A. Davis, Y. Zhang, R. Li, M. Salit, T. R. Gingeras and B. Oliver. 2011. Synthetic spike-in standards for RNA-seq experiments. Genome Res. 21(9): 1543–1551.

Li B, Dewey CN. 2011. RSEM: accurate transcript quantification from RNA-Seq data with or without a reference genome. BMC Bioinformatics. 12:323.

Li H, Durbin R. Fast and accurate short read alignment with Burrows-Wheeler transform. 2009. Bioinformatics. 25:1754–60.

Love MI, Huber W, Anders S. 2014. Moderated estimation of fold change and dispersion for RNA-seq data with DESeq2. Genome Biol. 15:550.

Quinlan AR, Hall IM. 2010. BEDTools: a flexible suite of utilities for comparing genomic features. Bioinformatics. 26:841–2.

Risso D, Ngai J, Speed TP, Dudoit S. 2014. Normalization of RNA-seq data using factor analysis of control genes or samples. Nat. Biotechnol. 32:896–902.

Ulitsky I, Bartel DP. 2013. lincRNAs: Genomics, Evolution, and Mechanisms. Cell. 154:26–46.

Zhao W, He X, Hoadley KA, Parker JS, Hayes DN, Perou CM. 2014. Comparison of RNA-Seq by poly (A) capture, ribosomal RNA depletion, and DNA microarray for expression profiling. BMC Genomics. 15:419.

